# Postsynaptic integration of excitatory and inhibitory signals based on an adaptive firing threshold

**DOI:** 10.64898/2026.03.26.714497

**Authors:** Oliver Gambrell, Pavol Bokes, Abhyudai Singh

## Abstract

A key component of intraneuronal communication is the modulation of postsynaptic firing frequencies by stochastic transmitter release from presynaptic neurons. The time interval between successive postsynaptic firings is called the inter-spike interval (ISI), and understanding its statistics is integral to neural information processing. We start with a model of an excitatory chemical synapse with postsynaptic neuron firing governed as per a classical integrate-and-fire model. Using a first-passage time framework, we derive exact analytical results for the ISI statistical moments, revealing parameter regimes driving precision in postsynaptic action potential timing. Next, we extended this analysis to include both an excitatory and an inhibitory presynaptic connection onto the same postsynaptic neuron. We consider both a fixed postsynaptic-firing threshold and a threshold that adapts based on the postsynaptic membrane potential history. Our analysis shows that the latter adaptive threshold can result in scenarios where increasing the inhibitory input frequency increases the postsynaptic firing frequency. Moreover, we characterize parameter regimes where ISI noise is hypo-exponential or hyperexponential based on its coefficient of variation being less than or higher than one, respectively.

## I. Introduction

The brain is a densely interconnected network of neurons that communicate through electrical signals known as action potentials (APs). The arrival of an AP at the active zone of a presynaptic axon terminal initiates the fusion of docked synaptic vesicles (SVs), which release neurotransmitters into the synaptic cleft. These neurotransmitters subsequently bind to receptors on the postsynaptic neuron and induce changes in the membrane potential level [1]. When the membrane potential is driven to a threshold potential, the postsynaptic neuron fires an AP. Following this, the membrane potential returns to a resting potential.

The time interval between successive APs, called the interspike interval (ISI), is of particular interest in neuroscience, as it encodes information about the rate, regularity of a neuron’s spike timing, and functionality within larger neuronal circuits. This functionality is predominantly controlled by the number of SVs that fuse upon an AP arrival, referred to as the quantal content (QC). QC and its effect on ISI have been extensively studied in prior works [2]–[7]. Specifically, the statistics of the number of docked SVs and QC have been modeled in terms of both fixed [4], [5], [8] and time-varying [9] presynaptic parameters. Models have also studied the effects of presynaptic regulation on SV dynamics, QC, and ISI [10]–[18]. As SV dynamics are inherently stochastic, stochastic models of QC and SV dynamics have been explored [19]–[25]. In our previous works, the distribution and moments of QC were studied in both the steady state [26] and transient [27] parameter regimes. In particular, analysis of experimental recordings in [27] shows that correlations between successive QCs are small in MNTB-LSO synapses, suggesting that simpler models could capture ISI statistics.

Motivated by these observations, we model QC as independent and identically distributed (i.i.d.) random variables and analytically derive the mean and noise of the ISI of a classical integrate-and-fire model utilizing a first-passage-time (FPT) [28]–[30] framework. Our analysis shows that ISI noise is minimized in two situations: at intermediate threshold levels and at intermediate QC levels. Interestingly, the minimum ISI decreases as the time constant increases.

After this, the model is extended to include both excitatory and inhibitory presynaptic neurons, which we call an EI network. Simulations of this model show that ISI noise is maximized at intermediate excitatory input frequencies for a fixed inhibitory input frequency when the threshold potential is fixed. Following this, we explore the effects of an adaptive threshold on the mean and noise in the ISI and find that the addition of presynaptic inhibition can increase the mean postsynaptic firing frequency. In particular, simulations show that the mean postsynaptic firing frequency increases at intermediate inhibitory input frequencies. In addition to the mean postsynaptic firing frequency, we also study the noise in the ISI, defined by its coefficient of variation. Our results on ISI noise demonstrate the existence of a critical inhibitory input frequency that partitions the inhibitory input frequency space into two regions: one where noise is hypo-exponential and the other where noise is hyper-exponential. We now begin discussing the synaptic model and then formulate its dynamics.

## II. Model formulation of synaptic transmission

In this section, we model the dynamics of an excitatory presynaptic neuron terminating onto a postsynaptic neuron. A schematic of the system is presented in Fig. 1-A. The model will now be mathematically formulated.

**Fig. 1.**
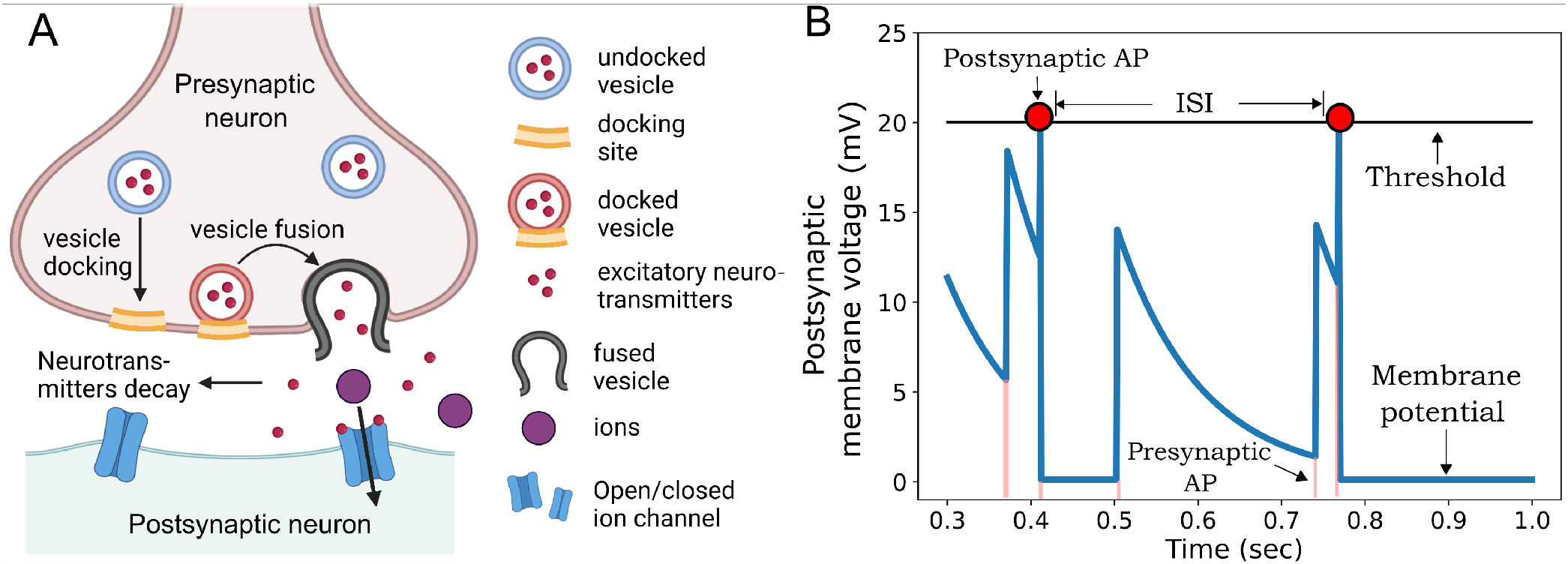
A schematic of synaptic transmission and a sample trajectory of the membrane potential from the associated synaptic model. **A**: Synaptic vesicles docked at docking sites at the active zone of the presynaptic axon terminal fuse with the membrane and release their neurotransmitter content into the synaptic cleft. These neurotransmitters then bind to receptors on the postsynaptic neuron, which facilitates the movement of positive ions between the inside and the outside of the postsynaptic neuron, changing its membrane potential. **B**: A simulated trajectory of the postsynaptic neuron’s membrane potential as modeled in equations (1) and (2). The membrane potential (blue line) depolarizes upon the arrival of presynaptic action potentials (red lines), which arrive via a Poisson process with rate *f*_*e*_. The number of synaptic vesicles (SVs) that fuse upon the arrival of an action potential (AP) follows a Poisson distribution with mean ⟨*b*_*e*_⟩. Between presynaptic APs, the membrane potential exponentially decays, with time constant *τ*_*v*_, to a resting potential, here 0. Once the membrane potential exceeds a threshold potential *v*_*th*_, the postsynaptic neuron fires an action potential (red circles). The ISI is the time between postsynaptic APs. Parameters: *f*_*e*_ = 10 *Hz*, ⟨*b*_*e*_⟩ = 15, *τ*_*v*_ = 100 *ms, v*_*th*_ = 20 *mV* .

We assume that presynaptic excitatory action potentials (APs) arrive via a Poisson process with rate *f*_*e*_. Upon the arrival of a presynaptic AP, synaptic vesicles (SVs) docked to docking sites on the presynaptic axon terminal fuse with the membrane wall. The fusion of SVs leads to the release of neurotransmitters into the synaptic cleft, which bind to ion channels on the postsynaptic neuron and induce a depolarization proportional to the number of synaptic vesicles that fuse. We model the process of SV fusion and subsequent postsynaptic depolarization with the following reset

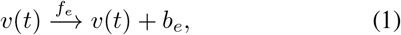

where *v*(*t*) represents the postsynaptic membrane potential, and *b*_*e*_ represents the number of SVs which fuse upon the arrival of a presynaptic AP. We refer to *b*_*e*_ as the quantal content (QC). For a given frequency of presynaptic stimulation, both the transient and steady-state QC have been shown to follow a binomial distribution in certain limits [26], [31]–[35]. Past analysis of mechanised stochastic models of SV docking/undocking and evoked release [27] has reported correlations between successive QC to be slightly anticorrelated. For modeling simplicity, we assume *b*_*e*_ to be an i.i.d random variable.

On the postsynaptic side, between excitatory presynaptic APs, the membrane potential returns to a baseline resting potential *v*_*r*_ via a classical leaky integrate-and-fire model as in

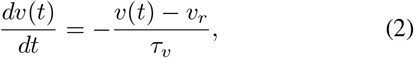

where *τ*_*v*_ is the positively valued membrane time constant that controls the rate at which the membrane potential returns to *v*_*r*_. We assume that *v*_*r*_ = 0.

If *v*(*t*) crosses a threshold potential *v*_*th*_, the postsynaptic neuron fires an AP and enters a refractory period where the membrane potential temporarily does not change in response to presynaptic APs. For simplicity, we assume that *v* instantly resets to *v*_*r*_ when *v* > *v*_*th*_. Our goal is now to quantify statistics of postsynaptic firing times.

The time interval between postsynaptic APs is called the inter-spike interval (ISI), which we define mathematically as

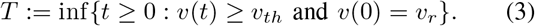

We study the mean ISI, ⟨*T* ⟩, and the noise in ISI, defined as

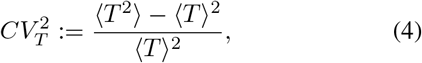

where ⟨ ⟩ denotes the expected value operator. Solving for the statistics of *T* is known as an FPT problem. FPT problems have been analyzed in great detail in cellular systems [28]–[30], lysis timing [36]–[39], gene expression [40]–[47], cellular event timing [48], [49], and previous works on neuronal modeling [22], [23], [50].

The FPT and its noise, when analyzed together, reveal information about neural coding and regularity. A schematic of the model and trajectories of *v*(*t*) are shown in Fig. 1-A and Fig. 1-B, respectively. Having formulated the synaptic model, we now formulate an expression for the moments of the ISI using an FPT framework.

## III. Exact analytical FPT derivation

In the previous section, we derived an i.i.d. model of a synapse and defined the FPT. Previous works have solved for the moments of the FPT using a small-noise approximation [20]–[22]. An exact analytical expression for the mean and noise of the FPT is now derived. The following analysis holds for any integer-valued distribution of *b*_*e*_.

### A. Survival function and moments

In order to derive the moments of the FPT, we define the survival function, which quantifies the probability of *v*(*t*) not crossing *v*_*th*_ as

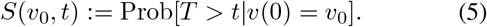

The survival function (5) depends the survival time *t* and the initial condition *v*_0_ = *v*(0) < *v*_*th*_ of the process. Although in this paper we are interested specifically in *v*_0_ = 0, the values at *v*_0_ > 0 are required to calculate the values of *v*_0_ = 0 using the backward equation approach adopted here.

The trivial equalities

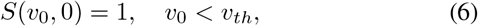

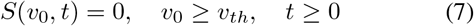

provide the initial and boundary conditions for (5).

We will be specifically interested in the FPT moments. The probability density function of the FPT is given by −∂*S*(*v*_0_, *t*)*/*∂*t*, so that the moments are defined by

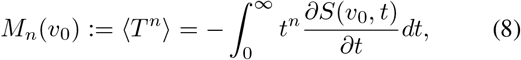

in which we make explicit the dependence of the FPT moments on the initial condition *v*_0_ = *v*(0) of the process. Substituting (7) into (8) yields

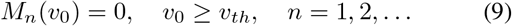

providing boundary conditions for the FPT moments.

### B. FPT threshold discontinuity

Since it takes at least one presynaptic AP to cross the threshold, the FPT is greater than or equal to the waiting time *T*_*nb*_ till the next presynaptic AP, which is exponentially distributed with mean 1*/f*_*e*_:

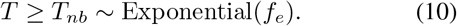

The inequality (10) implies that the FPT survival function must be bounded from below by the survival function of the exponential distribution:

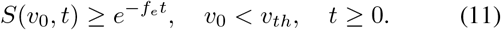

The FPT moments must also be greater than or equal to those of the exponential distribution:

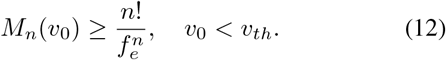

Combining (12) with the boundary condition (9), we obtain

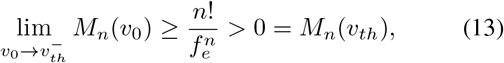

where the minus sign indicates that *v*_0_ approaches *v*_*th*_ from below. Thus, *M*_*n*_(*v*_0_) is discontinuous at *v*_0_ = *v*_*th*_.

We note that if the excitatory presynaptic AP frequency *f*_*e*_ is large, the lower bound (12) and the discontinuity height are small, which is consistent with a continuous FPT for diffusion processes [51].

### C. Delayed differential equation

Given the initial condition *v*_0_ ∈ (0, *v*_*th*_), the *n*-th FPT moment *f* (*v*_0_) = *M*_*n*_(*v*_0_) is obtained by solving the inhomogeneous linear delayed differential equation (see Appendix A for details)

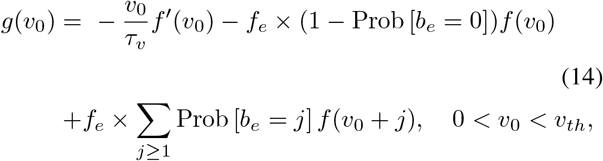

where *g*(*v*_0_) = − *nM*_*n*−1_(*v*_0_) and *f* (*v*_0_) = 0 for *v*_0_ ≥ *v*_*th*_. Equations (14) are solved iteratively for increasing moment order *n* ≥ 1, starting with *M*_0_(*v*_0_) = 1 for the 0-th moment.

While there are infinitely many solutions to (14) depending on the value *f* (*v*_*th*_), exactly one of them is convergent as *v*_0_ → 0 (see Appendix B). After finding the convergent solution, 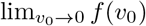 gives the desired FPT for the initial condition *v*(0) = 0.

Dependence of the exact FPT moments on model parameters is shown in Fig. 2-A,B,C. In Fig. 2-A, ISI mean and noise monotonically decrease as the input frequency increases. Fig. 2-B shows that the mean ISI monotonically decreases while the ISI noise depends non-monotonically on the mean QC. For Fig. 2-C, the ISI mean monotonically increases while the ISI noise depends non-monotonically on the threshold. Discontinuities in the ISI noise occur at integer values of the threshold, resulting in oscillations. In Fig. 2-B,C, ISI noise is exponential 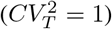 when ⟨*b*_*e*_⟩ and *v*_*th*_ are very small or very large. When these parameters are at intermediate values, noise is hypo-exponential 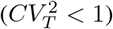.

**Fig. 2.**
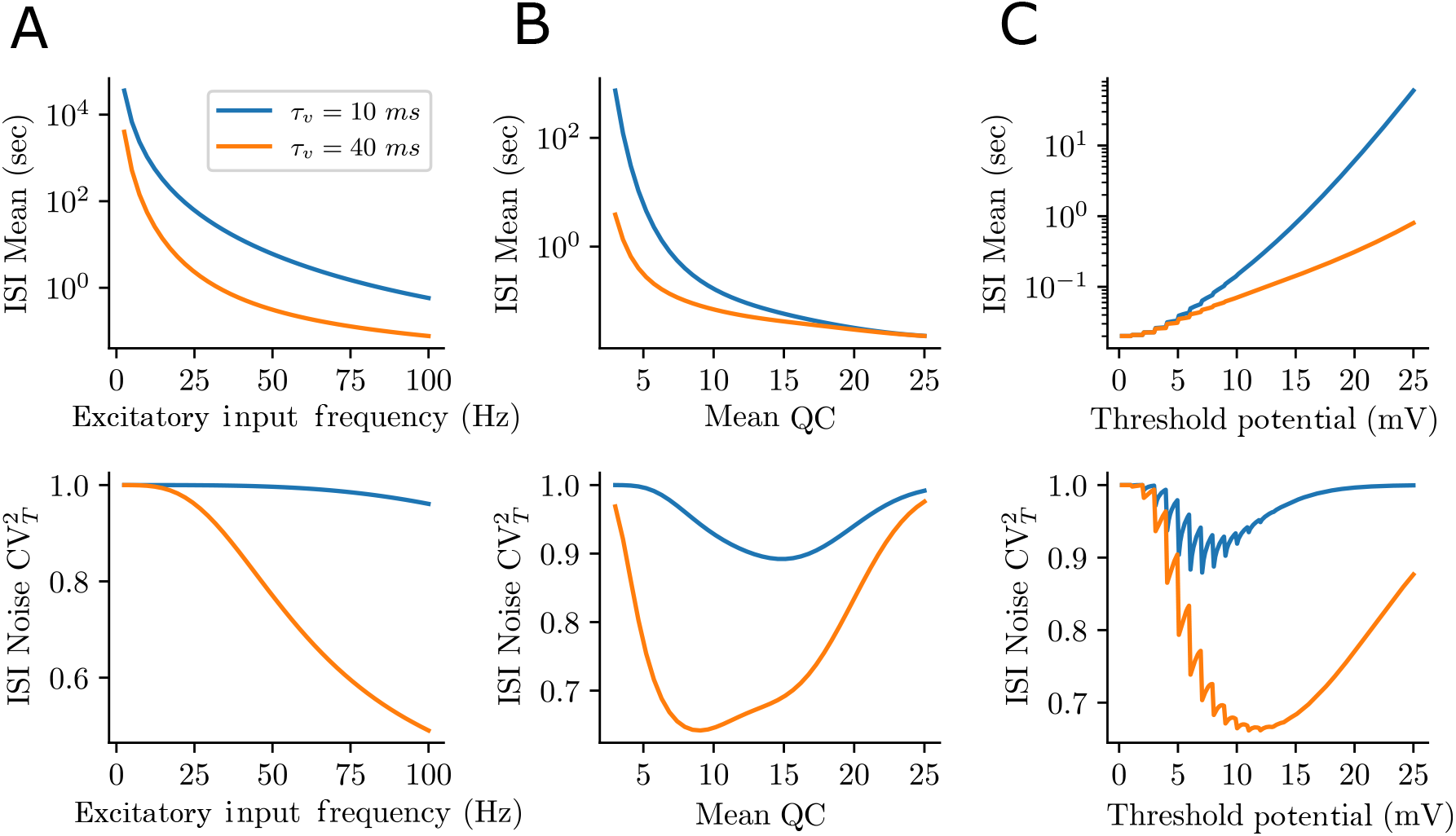
Characterizing the statistics of postsynaptic firing times as a function of both presynaptic and postsynaptic parameters. The synaptic model used for these figures is from equations (1) and (2), and the results of this figure are the solutions to equation (14). **A**: ISI mean (top) and ISI noise (bottom) as a function of the input frequency. Parameter values: *v*_*th*_ = 20 *mV*, ⟨*b*_*e*_⟩ = 5, *v*_0_ = 0 *mV* . **B**: ISI mean (top) and noise (bottom) as a function of the mean QC. Parameter values: *f*_*e*_ = 50 *Hz, v*_*th*_ = 20 *mV, v*_0_ = 0 *mV* . **C**: ISI mean (top) and noise (bottom) as a function of the threshold potential. Parameter values: *f*_*e*_ = 50 *Hz*, ⟨*b*_*e*_⟩ = 5, *v*_0_ = 0 *mV* .

Having formulated an expression for the moments of the ISI, in the following section, we expand the model by adding an inhibitory presynaptic neuron.

## IV. Model formulation for an excitatory-inhibitory (ei) neuronal network

In the previous section, we formulated a model for synaptic transmission of a single excitatory presynaptic input based on the classical integrate-and-fire model and derived its exact postsynaptic FPT moments. As many neuronal circuits receive inhibition, which is important in inter-spike interval (ISI) regulation [52]–[59], we extend the model by adding an inhibitory presynaptic input that terminates onto the postsynaptic neuron. In this extended model, we analyze the inverse of the mean ISI, called the mean postsynaptic firing frequency, due to its ease of interpretation. We refer to this model as the excitatory-inhibitory (EI) model.

The EI model schematic is presented in Fig. 3-A and is described as an excitatory and inhibitory presynaptic neuron independently firing APs onto the same postsynaptic neuron. The arrival times of both excitatory and inhibitory presynaptic APs follow independent Poisson processes with rates *f*_*e*_ and *f*_*i*_, respectively. The dynamics of the EI model are the same as in (1) and (2) with the addition of an inhibitory reset:

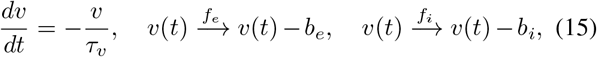

where *b*_*i*_ is the i.i.d. QC resulting from inhibitory presynaptic APs, *b*_*e*_ is the QC resulting form excitatory presynatpci APs, and *v* → *v*_*r*_ when *v* > *v*_*th*_.

**Fig. 3.**
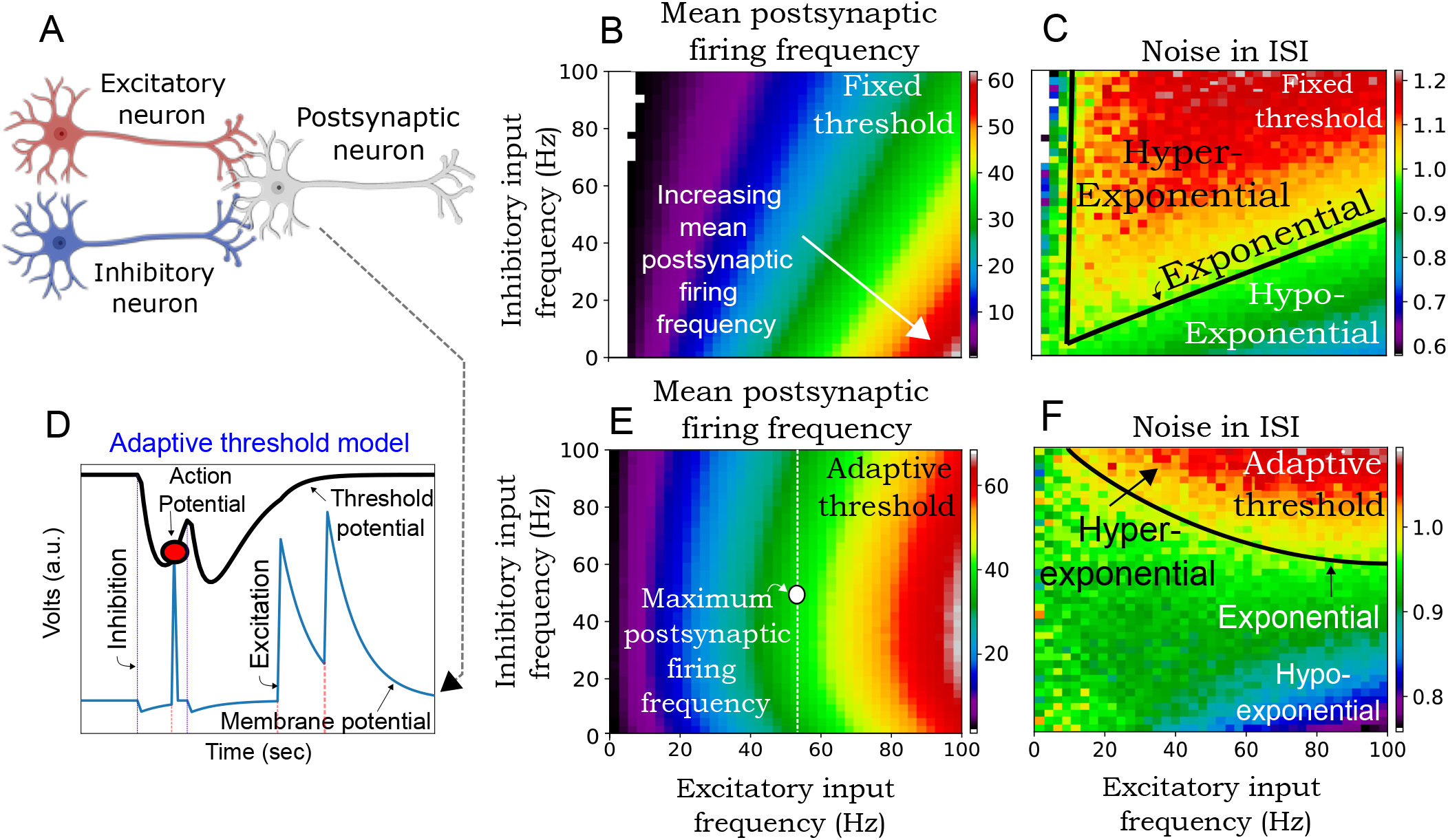
Inhibitory presynaptic APs can increase the mean postsynaptic firing frequency when the threshold potential adapts with recent firing history. The simulations in **B** and **C** come from the model in equation (15) and the simulations in **E** and **F** come from the adaptive threshold model in (16) and (17). In **B, C, E**, and **F**, excitatory and inhibitory QC follow gamma distributions with means ⟨*b*_*e*_⟩, ⟨*b*_*i*_⟩ and noise, 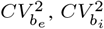, respectively. **A**: Excitatory-Inhibitory (EI) model schematic. The axon terminals of an excitatory and an inhibitory presynaptic neuron both terminate on the dendrites of a postsynaptic neuron. **B**: The mean postsynaptic firing frequency as a function of excitatory and inhibitory input frequency for the fixed threshold model. **C**: Noise in ISI for the fixed threshold model. **D**: Sample membrane potential and threshold potential trajectories for the adaptive threshold model. Presynaptic excitatory and inhibitory APs arrive via Poisson processes with rates *f*_*e*_ and *f*_*i*_ and either depolarize or polarize the membrane potential (blue line). When the membrane potential is driven below its resting potential by an inhibitory presynaptic AP (purple dotted line), the threshold potential (black line) evolves via equation (16) and (17). This facilitates postsynaptic AP firing (red circle) upon the arrival of excitatory presynaptic APs (red dotted line). **E**: Mean postsynaptic firing frequency for the adaptive threshold model. The white dashed line is the mean postsynaptic firing frequency at 53 *Hz* for a varying inhibitory input frequency. The white circle shows that the maximum postsynaptic firing frequency occurs at an inhibitory input frequency of 55 *Hz*. **F**: Noise in ISI for the adaptive threshold model. Parameters: *v*_*th*_ = 20 *mV*, ⟨*b*_*e*_⟩ = 20 *mV*, 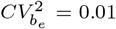, ⟨*b*_*i*_⟩ = 20 *mV*, 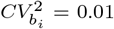, *τ*_*v*_ = 10 *ms, v*_*b*_ = 20 *mV, v*_*low*_ = 10 *mV, v*_1_ = −1 *mV* .

In Fig. 3-B, and Fig. 3-C, we plot the mean postsynaptic firing frequency and ISI noise of the EI model as a function of both the excitatory and inhibitory input frequency. From Fig. 3-B, we see that the mean postsynaptic firing frequency becomes increasingly sensitive to changes in the inhibitory input frequency with increasing excitatory input frequency.

Recall that a hypo-exponential, exponential, hyper-exponential random variable has its coefficient of variation < 1, 1, and > 1, respectively. Fig. 3-C, the excitatory-inhibitory input frequency space is partitioned into regions where ISI noise is hypo-exponential, exponential, and hyper-exponential. Interestingly, for an inhibitory input frequency over 4 *Hz*, noise is maximized at intermediate excitatory frequencies. Additionally, this maximum noise level increases with increasing inhibitory input frequency.

In this section, we analyzed the EI model for a fixed threshold potential. In the following section, the threshold potential will adapt based on the previous AP firing history.

## V. Excitatory-inhibitory neuronal network with an adaptive threshold

In previous sections, we analyzed ISI statistics for an EI network and simulated the ISI and its noise for a fixed threshold. This assumption, however, fails to capture the probabilistic nature of ion channel conductances such as sodium and potassium [60]. Previous works attempted to model these effects by incorporating an adaptive threshold [61]–[66], which improved neuronal models [67], [68]. With this understanding, we now modify our EI model by allowing the threshold potential to change in response to postsynaptic activity. In particular, the effects of post-inhibitory facilitation are modeled as observed in [69], which is characterized by the facilitation of postsynaptic spiking following brief presynaptic inhibitory APs.

In order to incorporate the facilitation of postsynaptic spiking into our model, the threshold potential is now a function of *v*(*t*) that evolves via

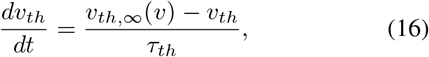

where *τ*_*th*_ is the threshold time constant that controls the rate at which the threshold potential approaches a steadystate level *v*_*th*,∞_(*v*), defined as

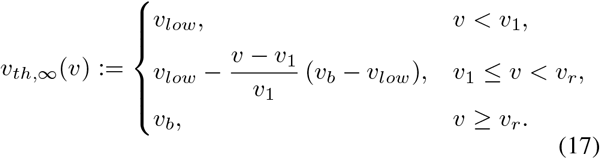

In the case of depolarization (*v* > *v*_*r*_), the threshold is at a fixed value (as before). But for the hyperpolarized case (*v* < *v*_*r*_), the threshold potential decreases with decreasing *v* to a basal level *v*_*low*_ which occurs when *v* = *v*_1_. Here we assume this decrease to follow a piecewise linear form as shown in equation (17). A sample trajectory of the adaptive threshold model is plotted in Fig. 3-D, which shows a postsynaptic AP occurring when *v*_*th*_ < *v*_*b*_.

In Fig. 3-E and Fig. 3-F, we simulate the inverse of the mean ISI, called the mean postsynaptic firing frequency, and the noise 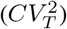 in the ISI as a function of both the excitatory and inhibitory input frequency, respectively. From Fig. 3-E, we see that the mean postsynaptic firing frequency is maximized at intermediate inhibitory input frequencies for a fixed excitatory input frequency. Additionally, this maximum value increases with the excitatory input frequency. The ISI noise plot in Fig. 3-F shows that the excitatory-inhibitory input frequency space is partitioned into hypo-exponential 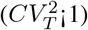, exponential 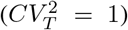, and hyper-exponential 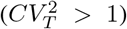 regions similar to the fixed threshold model of Fig. 3-C. In contrast to Fig. 3-C, there exists a critical inhibitory input frequency which partitions the inhibitory input frequency space into two regions: one where noise is always hypo-exponential and one where noise can become hyper-exponential with increasing excitatory input frequency. Furthermore, simulations reveal the existence of a critical inhibitory frequency. For frequencies below this level, noise is always hypo-exponential, and for frequencies above this level, noise is capable of transitioning from hypo to hyper-exponential with increasing excitatory input frequency. Additionally, as the inhibitory input frequency increases above this critical level, noise transitions from hypo to hyper-exponential at lower excitatory input frequencies.

## VI. Conclusion

This work explored the postsynaptic inter-spike interval (ISI) statistics of a leaky integrate-and-fire model (Fig. 1). For this model, quantal content (QC) was assumed to be independent and identically distributed (i.i.d.) based on previous works showing low correlations between successive QC [27]. The key contribution of this paper was the derivation of an analytical expression for the moments of the ISI using a first-passage time (FPT) framework. Our analysis revealed that noise is always hypo-exponential (coefficient of variation is less than one) in this model (Fig. 2), implying that the postsynaptic firing frequency exhibits regularity.

Following this, we extended our model by adding an inhibitory presynaptic connection onto the same postsynaptic neuron and simulated the mean postsynaptic firing frequency and ISI noise (Fig. 3-A). The analysis was done for both a fixed threshold and a threshold that adapts based on the postsynaptic membrane potential history. Our results for the case of a fixed and adaptive threshold showed the existence of a parameter space where noise is hyper-exponential. In the adaptive threshold model (Fig. 3-D), the parameter space where noise is hypo-exponential is enlarged when compared to the fixed threshold model due to the regulatory effect of adaptive thresholds on postsynaptic firing frequencies [70].

Similar to our findings in Fig. 2, recent studies on intracellular event timing using the bacteriophage lysis system have shown experimentally that timing noise varies non-monotonically with lysis threshold and noise is minimized at an optimal threshold [71].

In future work, we will derive analytical expressions for the ISI of the EI model for different types of adaptive thresholds. Specifically, the threshold potential will model regulatory mechanisms as in [70], and increase following postsynaptic APs. Additionally, models receiving solely inhibition and no excitation will be studied, as experimental evidence has shown that inhibition by itself can generate postsynaptic APs [72].

## Appendix A: Backward equations

The ISI survival function *S*(*v*_0_, *t*) and moments *M*_*n*_(*v*_0_) are determined for all initial conditions 0 ≤ *v*_0_ < *v*_*th*_ by solving specific backward equations. The equations are based on the backward operator ℒ, which takes a function *f* (*v*_0_) of a positive real variable *v*_0_ and returns a new function

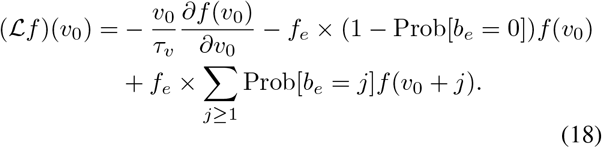

The backward operator is the sum of a differential operator (due to deterministic decay) and a difference operator (due to random bursts drawn from a discrete distribution). We note that for continuous burst distributions, the difference operator needs to be replaced by an integral operator, a situation we do not discuss here.

The ISI survival probability *S*(*v*_0_, *t*) satisfies the backward equation [51]

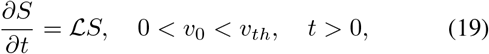

which is subject to the initial and boundary conditions (6)– (7). Multiplying (19) by powers of *t*, integrating, and using (8), we obtain backward equations for the ISI moments:

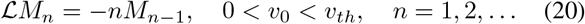

which are subject to the boundary condition (9). Setting (18) into (20) yields (14).

## Appendix B: Existence and uniqueness result

Below, we define solution spaces and formulate the uniqueness and existence result for (14). The function space Ω_0_ is the one in which the ISI moment is sought. However, it is also useful to consider a broader space of functions Ω that allow solutions that are divergent as *v*_0_ → 0.

### Definition 1

(Solution spaces for (14)). *For a function f* : (0, ∞) → ℝ, *we write f* ∈ Ω *if assumptions (A1)–(A3) are satisfied; we write f* ∈ Ω_0_ *if assumption (A4) is additionally satisfied:*

*(A1) f* (*v*_0_) = 0 *for v*_0_ ≥ *v*_*th*_ *(the boundary condition)*

*(A2) f* (*v*_0_) *is continuous for v*_0_ ∈ (0, *v*_*th*_) *with a finite onesided limit* 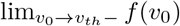

*(A3) f* ^*′*^(*v*_0_) *exists and is continuous for v*_0_ ∈ (0, *v*_*th*_) \ {*v*_*th*_ − *j* : *j* ∈ ℕ}. *For v*_0_ ∈ {*v*_*th*_ − *j* : *j* ∈ ℕ, *v*_*th*_ − *j* > 0}, *finite one-sided limits* 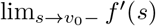 (*s*) *and* 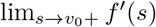 *exist*

*(A4) A finite one-sided limit* 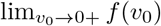 *exists*

In the next section, we prove the following theorem. The proof is constructive, leading to a method for enumerating ISI moments.

### Theorem 1

(Existence and uniqueness result). *For g* ∈ Ω_0_, *there exists a unique solution f* ∈ Ω_0_ *to the delayed differential equation* (14).

*Proof:* Since ℒ is linear and first-order, a solution *f* (*v*_0_) to (14) can be decomposed into the homogeneous and particular parts

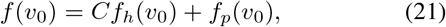

where *f*_*h*_(*v*_0_) and *f*_*p*_(*v*_0_) satisfy

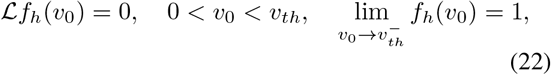

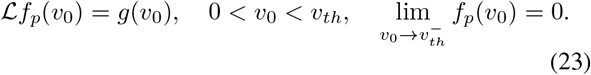

Indeed, adding up *C* times (22) to (23) implies that (21) satisfies (14); we also have 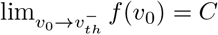

Solutions (22) and (23) are initialised with the left-limit value at the threshold *v*_0_ = *v*_*th*_ and integrated numerically in the direction of decreasing *v*_0_. It is clear, by construction, that the solutions satisfy (A1)–(A3), i.e. *f*_*h*_, *f*_*p*_ ∈ Ω. Therefore, their linear combination (21) also satisfies *f* ∈ Ω.

What remains is to ascertain whether *f* ∈ Ω_0_, i.e., whether (A4) is satisfied. The key to this is the asymptotic behaviour as *v*_0_ → 0^+^ of solutions to (14). Seeking dominant balance, we equate the first two terms 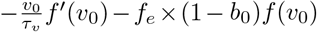 to zero and solve in *f*, obtaining the power law asymptotics

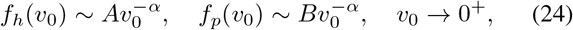

where *A, B* ≠ 0 and

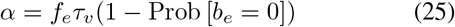

*α* = *f*_*e*_*τ*_*v*_(1 − Prob [*b*_*e*_ = 0]) (25) gives a normalised rate of non-zero bursts. Note that *f*_*h*_, *f*_*p*_ ∉ Ω_0_, as they fail to meet assumption (A4). By (21) and (24),

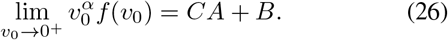

In order to meet (A4), we have to set

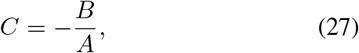

thus ensuring that *f* ∈ Ω_0_. Clearly, condition (27) determines the solution uniquely, hence Theorem 1 is proved.

